# Over 50000 metagenomically assembled draft genomes for the human oral microbiome reveal new taxa

**DOI:** 10.1101/820365

**Authors:** Jie Zhu, Liu Tian, Peishan Chen, Mo Han, Liju Song, Xin Tong, Zhipeng Lin, Xing Liu, Chuan Liu, Xiaohan Wang, Yuxiang Lin, Kaiye Cai, Yong Hou, Xun Xu, Huanming Yang, Jian Wang, Karsten Kristiansen, Liang Xiao, Tao Zhang, Huijue Jia, Zhuye Jie

## Abstract

The oral cavity of each person is home for hundreds of bacterial species. While taxa for oral diseases have been well studied using culture-based as well as amplicon sequencing methods, metagenomic and genomic information remain scarce compared to the fecal microbiome. Here we provide metagenomic shotgun data for 3346 oral metagenomics samples, and together with 808 published samples, assemble 56,213 metagenome-assembled genomes (MAGs). 64% of the 3,589 species-level genome bins contained no publicly available genomes, others with only a handful. The resulting genome collection is representative of samples around the world and across physiological conditions, contained many genomes from Candidate phyla radiation (CPR) which lack monoculture, and enabled discovery of new taxa such as a family within the Acholeplasmataceae order. New biomarkers were identified for rheumatoid arthritis or colorectal cancer, which would be more convenient than fecal samples. The large number of metagenomic samples also allowed assembly of many strains from important oral taxa such as *Porphyromonas* and *Neisseria*. Predicted functions enrich in drug metabolism and small molecule synthesis. Thus, these data lay down a genomic framework for future inquiries of the human oral microbiome.

---

The human microbiome has been implicated in a growing number of diseases. The majority of microbial cells is believed to reside in the large intestine^1^ and cohorts with fecal metagenomic data contain over 1000 individuals^2, 3^. For the oral microbiome, hundreds of metagenomic shotgun-sequenced samples have been available from the Human Microbiome Project (HMP) and for rheumatoid arthritis^4–6^. A number of other diseases studied by Metagenome-wide association studies (MWAS) using gut microbiome data also indicated potential contribution from the oral microbiome in disease etiology^7–12^. Although the MWAS on rheumatoid arthritis was based on a *de novo* assembled reference gene catalog for the oral microbiome^6^, analyses on bacterial genomes would be more desirable. And when oral samples show comparable or better sensitivity and accuracy for disease diagnosis, prognosis or patient stratification than fecal samples, oral samples would be much more convenient as they could be available at any time and taken at a fully controlled setting witnessed by trained professionals. Unlike the anaerobic environment for the gut microbiome, the oral microbiome is believed to be well covered by culturing^13^, and analyses by 16S rRNA gene amplicon sequencing or polymerase chain reaction (PCR) are common. Recently published large-scale metagenomic assembly efforts mostly included fecal metagenomic data^14–16^. It is not clear how much is really missing for the oral microbiome. The saliva, in particular, seems to have more bacterial species per individual than the fecal microbiome^17^.

After getting contigs using assembly algorithms suitable for metagenomic data^18^, a central idea used by metagenomic binning algorithms is that genes or contigs that co-vary in abundance among many samples belong to the same microbial genome^8, 19–21^. Large cohorts are therefore prerequisites for high-quality assembly.

Here we present 3346 new oral metagenomic samples, and 56,213 metagenome-assembled genomes (MAGs) which represent 3,589 species-level clades, revealing new taxa as well as substantially complementing the genomic content of known species. This genome reference are highly representative of metagenomic samples not used in assembly, and could facilitating culturing, functional screens as well as disease diagnosis and modulation based on the oral microbiome.

## RESULTS

### Draft genomes assembled from oral metagenomic data

In order to substantially increase the amount of oral microbiome data, we shotgun sequenced 2284 saliva and 391 tongue dorsum samples from the 4D-SZ cohort^3, 12, 22^, 671 saliva samples from five ethnic groups of Yunnan province, producing over 43.19 terabytes of sequence data (**Supplementary Table 1**). Together with 808 published samples from 5 studies^6, 23–26^ that have not been used for metagenomic assembly (**Supplementary Table 1**), a total of 4,154 oral samples with metagenomic data were obtained. The data in each sample was single assembled into contigs using SPAdes^27, 28^(Fig. 1a, **Supplementary Table 1**). Binning was then performed using MetaBAT2^21^ for the 39,458,119 contigs longer than 1.5kb, leading to 56,213 metagenome-assembled genomes (MAGs), 15,013 of which were of high quality according to recently agreed standards^29^ using CheckM^30^ (>90% completion, <5% contamination, Fig. 1a). The remaining 41,200 also reached the standards for medium-quality MAGs (>50% completion, <10% contamination), while low-quality assemblies were not further analyzed (**Supplementary Table 2**). The median constructed MAGs per sample is 12, with highest from ZhangX_2015 (Supplementary Figure 1a).

**Figure 1.**
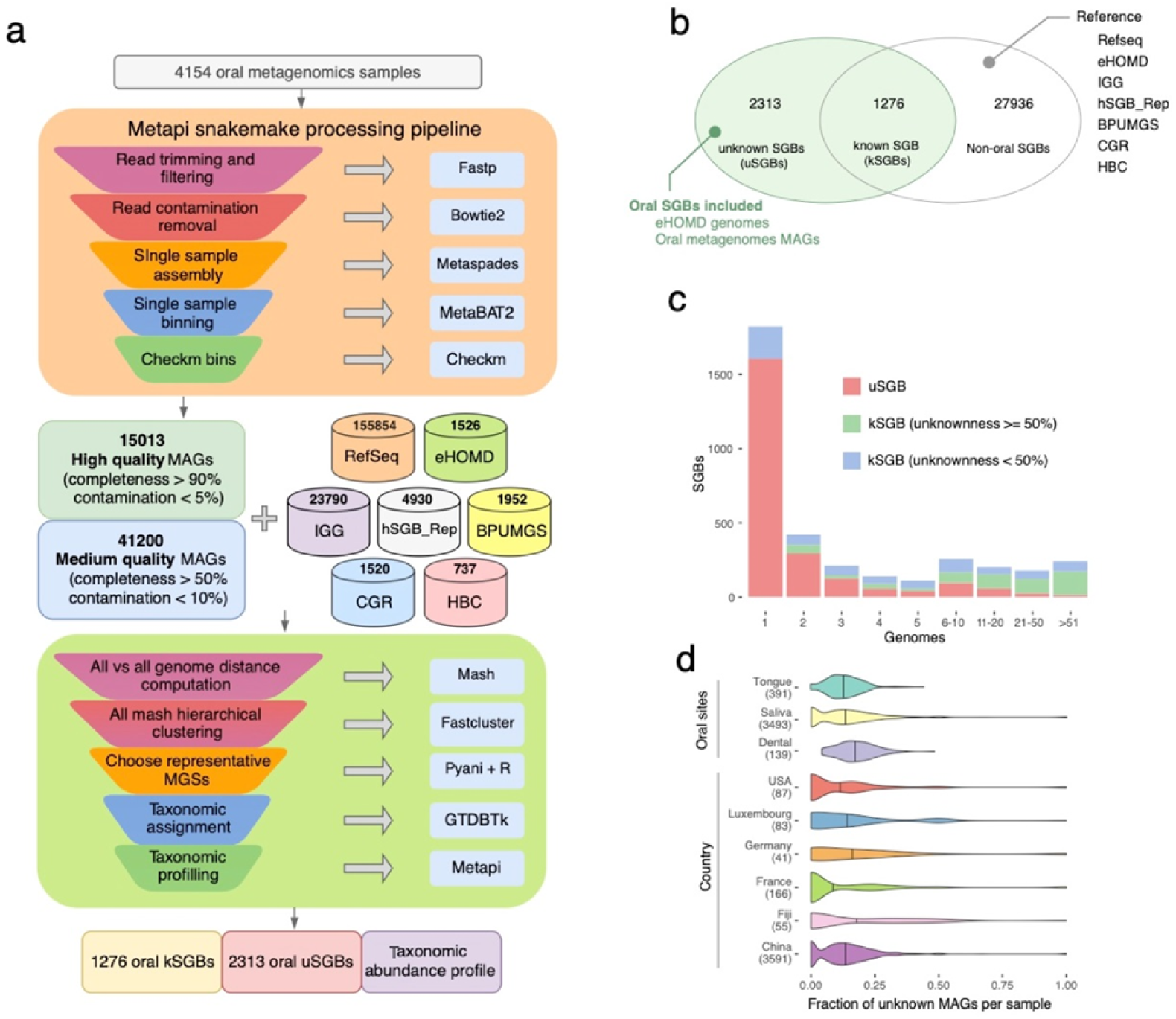
3,589 oral SGBs assembled from 4154 (3346 new sequence) meta-analyzed oral-wide metagenomes. **a**, Species genome bins construction workflow. **b**, Overlap of oral assembled genomes and reference genomes. kSGBs contains both existing microbial genomes (including other metagenomic assemblies) and genomes reconstructed here. uSGBs are only genomes reconstructed here and without existing isolate or metagenomically assembled genomes. Non-oral SGBs contains kSGBs that are not sourced from human oral samples. **c**, Genome numbers distribution of uSGBs and kSGBs. **d**, Distribution of the fraction of uMAGs in each sample by oral sites and country.

### New genomes from the new samples

To evaluate the novelty of the assembled genomes, delineate their taxonomy and potential source, we comprehensively incorporated 190,309 existing isolate or metagenome-assembled genomes from NCBI Refseq, eHOMD^31^ (the expanded Human Oral Database), and recent publications^14–16, 32^ including our culture collection from fecal samples in Shenzhen^33^ (Fig. 1a).

Species-level clusters (SGBs, for species-level genome bins) were computed for the over 0.25 million genomes following multiple steps (Fig. 1a, see Methods for details), defined as at least 95% average nucleotide identity (ANI) and at least 30% overlap of the aligned genomes. The clustering well-collapsed the genomes, with about 10-fold reduction in number, i.e. resulting in around thirty thousand species. Besides the 27,936 species that were non-oral according to reference genomes in the cluster (defined in Fig. 1b), 2,313 clusters (64% of the total oral species) only contained our MAGs (denoted uSGBs for unknown SGBs), some of which were repeatedly captured in our data, with more than 50 genomes each (Fig. 1b,c); the 1,276 known oral SGBs (kSGBs) could be further divided according to the percentage of reference genomes in the cluster. Interestingly, kSGBs with over 50% unknown genomes outnumbered kSGBs with 0-50% unknown genomes for clusters containing 10 or more genomes (Fig. 1b,c), underscoring the discovery power of large metagenomic cohorts. And the top three contributions of uSGBs are 4D_SZ (1441), ZhangX_2015 (445) and Yunnan (334) (Supplementary Figure 1b). Comparing the ratio of new MAGs in the samples, we retrieved a greater fraction of previously unknown genomes in dental compared to saliva or tongue samples, even though we did not take dental samples for this large cohort (Fig. 1d). This ratio also appeared to differ between cohorts, with less than 10% unknown for samples from France or the U.S., and more newly matched uSGBs for samples from Fiji, Germany and Luxemburg (Fig. 1d). The large cohort available from this study is crucial for the retrieval of novel oral species, contributing over 2000 uSGBs, greatly expanding our knowledge of oral microbiome diversity.

### Close to 90% representation of oral metagenomics data by the genomes

We next examined the ability of this species-level genomes set to represent metagenomic shotgun data. We assessed the percentage of reads that could align to cultured genomes only (eHOMD) and cultured complemented by metagenomically assembled genomes. The median was 66.99% mapping with the 1526 genomes from eHOMD (Fig. 2a). The 4930 representative human SGBs from a recent large-scale assembly study that included available oral metagenomic samples^16^ led to 79.72% mapping, and the representative oral 3589 SGBs from the current study instead led to 88.06% mapping (median for all samples), especially for metagenomes from the U.S. and Germany; and a median of 85.29% mapping even for 81 saliva and subgingival metagenomes from three cohorts that were not used in the assembly process^34–36^ (Fig. 2a**, Supplementary Table 1**). Across physiological states, our SGBs well represented pregnant samples from the U.S. (reaching 92.99% mapping), RA (reaching 90.69% mapping) and diabetes (reaching 83.99% mapping) (Fig. 2b). Such a high degree of representation of metagenomic data across geography, ethnicity, age and physiological states suggest that the expanded genomic content of oral SGBs could serve as a starting point for quantitive taxonomic and functional analyses of the human oral microbiome.

**Figure 2.**
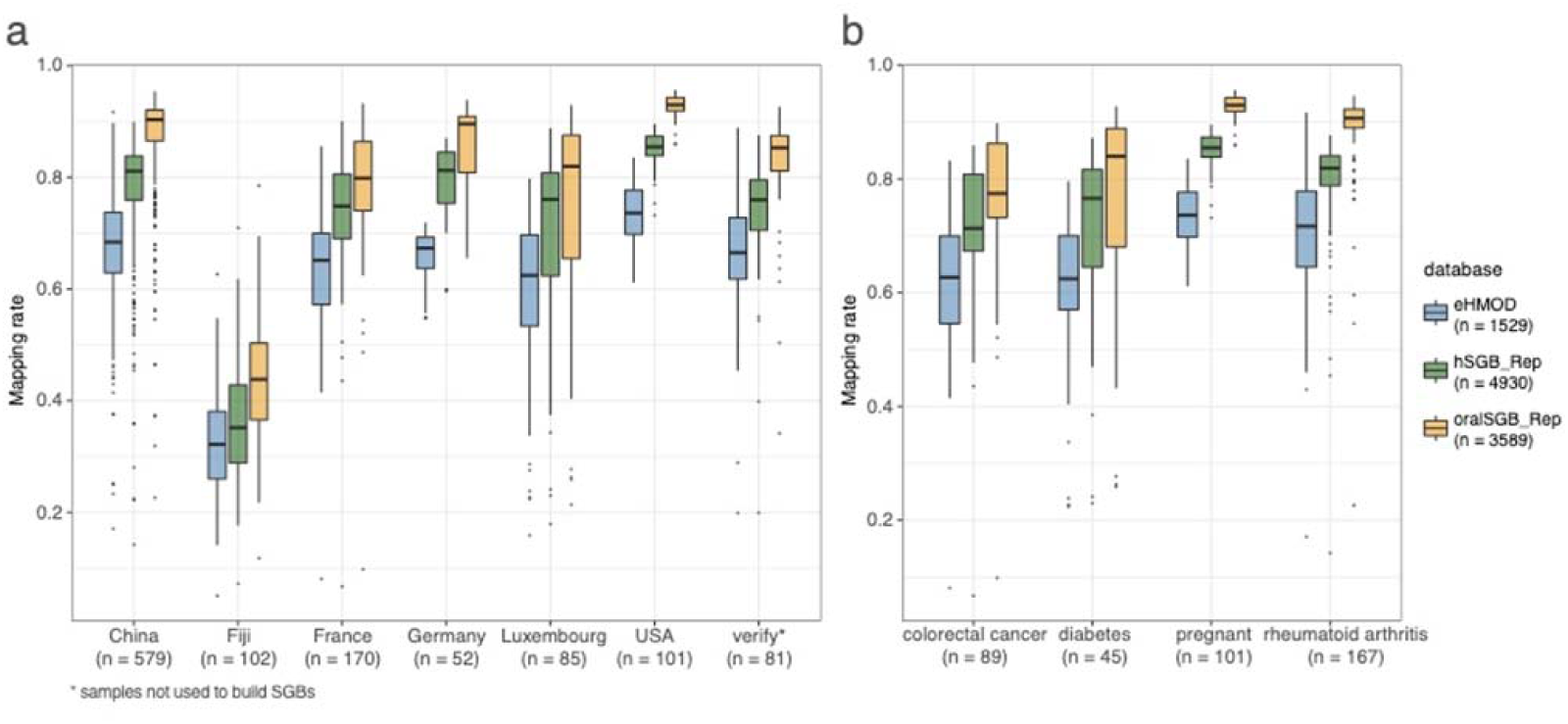
The expanded genome set substantially increases the mappability of oral metagenomes. The raw reads from all the 808 public samples, 100 subsampled of 2674 China Shenzhen and 671 China Yunnan, and 81 additional verify samples which not used to build SGBs were mapped against eHOMD, representative human SGBs (hSGB_Rep) and representative oral SGBs (oralSGB_Rep). Among three databases, our representative oral SGBs have the highest raw-read mappability in all country and verify datasets.

### Taxonomic landscape of the oral microbial genomes

We constructed a phylogenetic tree for the 3,589 oral SGBs, and similar to the gut microbiome, *Firmicutes* took up the largest number of branches (1248 clusters, 12307 genomes, Fig. 3c). The other 46276 genomes distributed into 15 phyla, including major human oral phyla such as *Actinobacteria* (490 SGBs, 6477 genomes), *Bacteroidetes* (368 SGBs, 23409 genomes), *Proteobacteria* (364 SGBs, 7570 genomes), *Campylobacteriota* (280 SGBs, 1841 genomes), and *Fusobacteriota* (145 SGBs, 1998 genomes) (Fig. 3, **Supplementary Table 3**). uSGB accounted for 181.27% increase in the reconstructed phylogenetic branch length, with over 80% of the diversity in *Campylobacterota* phylum contributed by the new uSGBs, follow by over 70% for *Patescibacteria* and *Fusobacteriota* (Fig. 3b), which seemed overlooked by culturing studies. We estimated there is median of 210 SGBs with the relative abundance higher than 0.001 per sample (Supplementary Figure 1c). Besides uSGB are also very high abundance, explained for 68.10% of richness and 65.23% of relative abundance per sample (Supplementary Figure 1d,e). Our MAGs greatly expanded the species or strains diversity within each phylum. As many as 596 SGBs from 4006 genomes belonged to the candidate superphylum of *Patescibacteria* (*Parcubacteria*, also known as OD1), which only have 157 kSGB with 3115 reference genomes. We note a few not so well studied phyla that were interesting in analogy to the gut microbiome. *Akkermansia* is the only genus from Verrucomicrobiota in the human gut and intensively pursued for its role in health and diseases, and Verrucomicrobiota and Spirochaetota take up a greater fraction in Hadza hunter gatherers compared to developed countries^37^. Here we identified 6 genomes in 3 SGBs for Verrucomicrobiota, and 900 genomes in 67 SGBs for Spirochaetota. 121 reference genome was only available for 32 SGB within Spirochaetota. 198 SGBs with 1169 genomes belong to the candidate division Saccharibacteria (TM7) (Fig. 3a, **Supplementary Table 3**).

**Figure 3.**
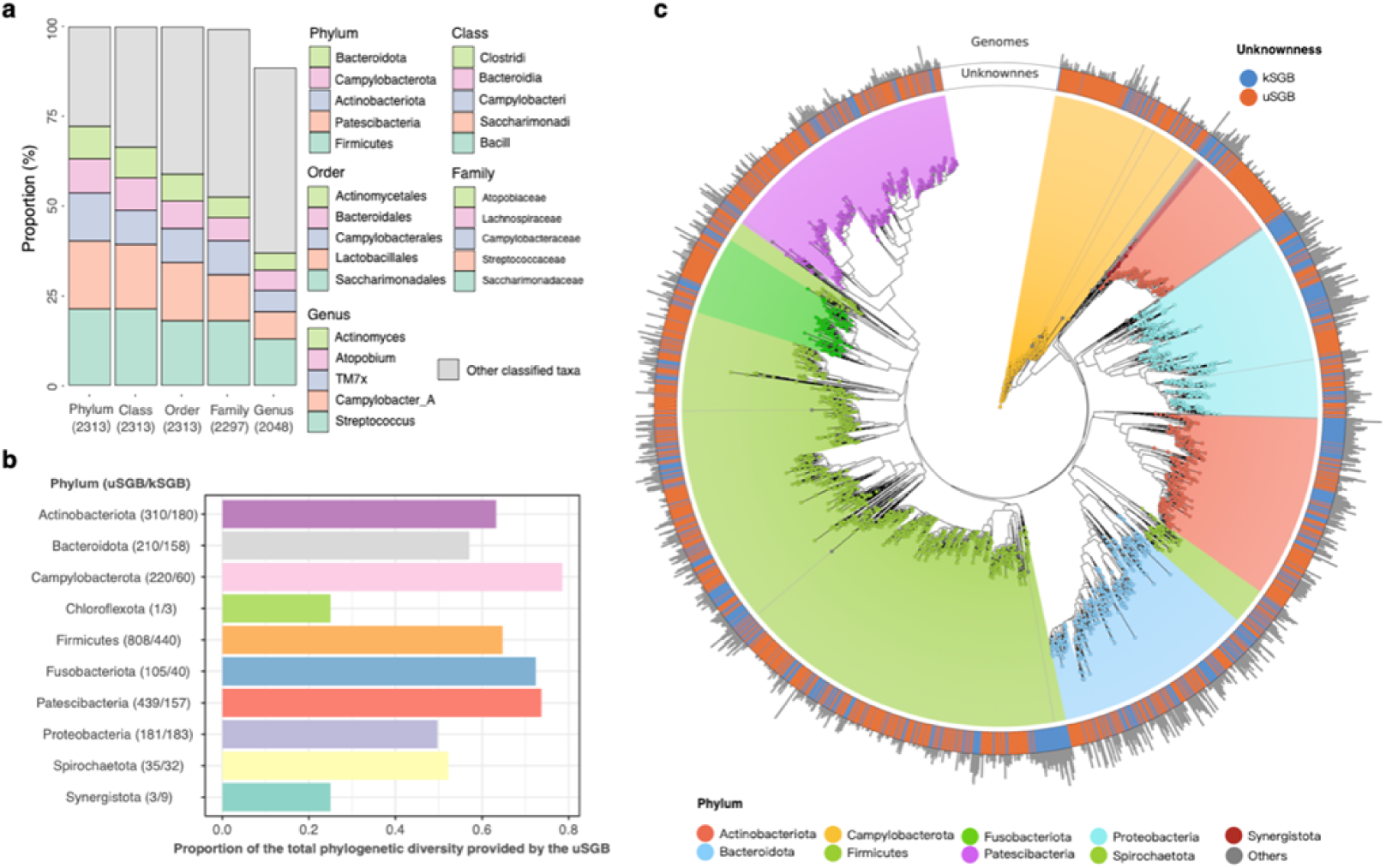
Phylogeny of representative oral SGBs. **a**, Taxonomic composition of the 2,313 uSGB, Only the five most frequently observed taxa are shown in the legend, with the remaining lineages grouped as “other classified taxa”. **b**, Proportion of the total phylogenetic diversity provided by the uSGB. **c**, oral-associated microbial phylogenetic tree of representative genomes from 3589 species-level genome bin (SGB).

At the genera level, *Streptococcus*(460 SGBs), *Campylobacter*(279 SGBs), *Actinomyces*(184 SGBs), *Prevotella*(159 SGBs), *Atopobium*(146 SGBs) were the major genera in the SGBs(**Supplementary Table 3**). 265 of the 2313 uSGBs had taxonomic information until order or family, but cannot be annotated to a known genus. The top three uSGB classified families were Saccharimonadaceae (17.99%), Streptococcaceae (12.88%) and Campylobacteraceae (9.51%), whereas the most assigned genera were Streptococcus (12.88%), Campylobacter_A (7.65%) and TM7x (5.92%) (Fig. 3c).

### A new family with small genomes

In the Acholeplasmatales order (Mollicutes class) of the *Tenericutes* phylum, a number of our uSGBs with high-quality MAGs formed a clade distinct from *Acheloplasma* and *Candidatus Phytoplasma*, with shallow branches within the clade (Fig. 4a). The genome size of this genome-defined family, which we temporarily denoted as *Ca. Bgiplasma*, is 0.69±0.05 Mbp, which is similar to *Candidatus Phytoplasma* (0.64±0.14 Mbp), but much smaller than *Acheloplasma* (1.50±0.20 Mbp). Genomes of such small size were discarded in early efforts of metagenomic assembly^19^, but we now know *Ca. Bgiplasma* are complete entities according to single-copy marker genes in CheckM (**Supplementary Table 2**). The GC content of the three clades were also different. *Ca.* Bgiplasma family was more towards normal GC content (34.57±0.21%), not as low as *Acheloplasma* (30.99±1.75%) and *Candidatus Phytoplasma* (25.98±2.68%) (**Supplementary Table 5**). Despite the lack of deep branches, the ANI distribution of uSGB within *Ca. Bgiplasma* family showed two separate groups at genus-level divergence (ANI <85%) (Fig. 4b), illustrating diversity within this new family. This 11 uSGBs comprising 29 MAGs contribute more than 0.1% relative abundance in 209 samples, indicating that this family is an potentially important but so far uncharacterized clade in the oral microbiome.

**Figure 4.**
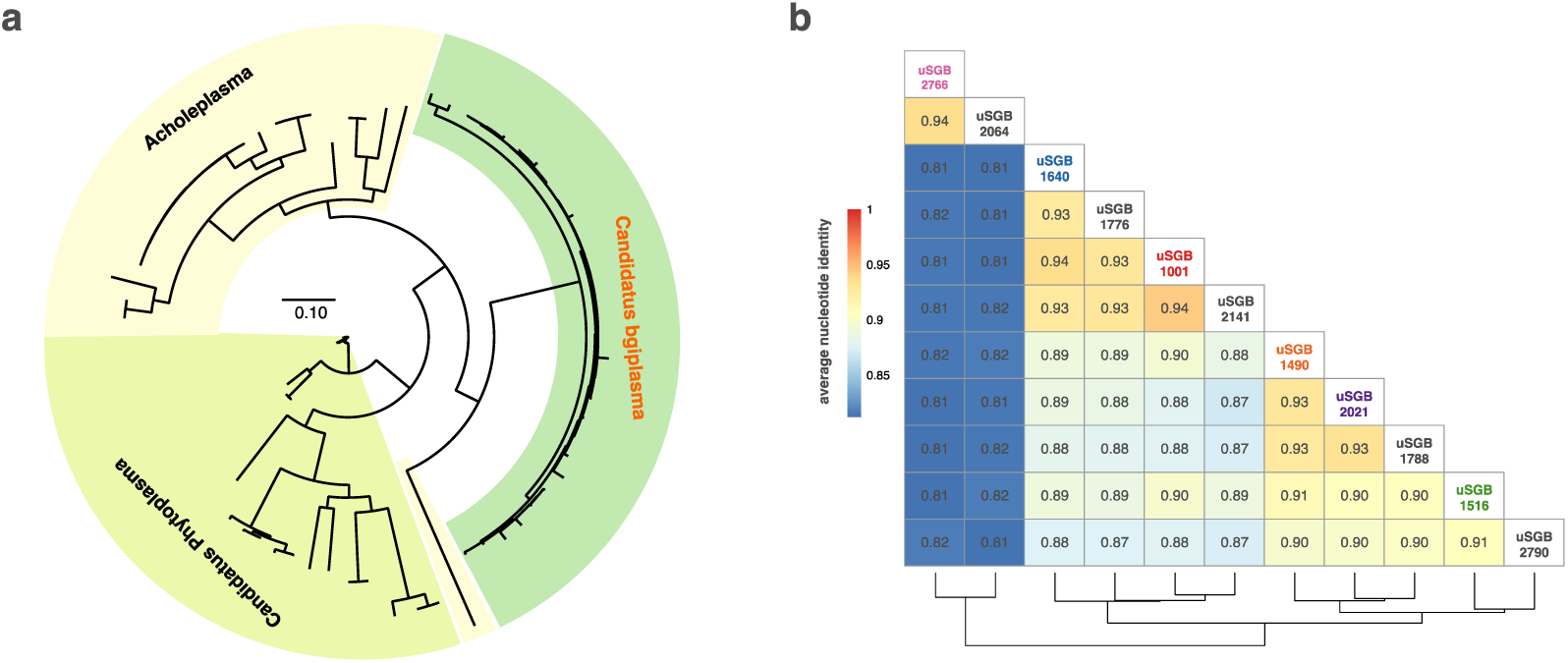
A new candidatus family is found within the Acholeplasmataceae Order. **a**, phylogenetic tree from all MAGs in the new candidatus family(Candidatus bgiplasma) and known genomes in Acholeplasmataceae Order. Supplementary Figure 2a reports the detail of phylogenetic tree in Candidatus bgiplasma. **b**, Average nucleotide identity(ANI) between all uSGB in the *Ca.* bgiplasma represent clearly two genus clades which ANI less than 0.85. Unknown SGBs without HQ MAGs are left black.

5.53 M genes (87.97% of total) of representatives genome of SGBs can be annotated by EggNOG mapper^38, 39^ with the rate of annotation 89.55% for uSGBs and 81.83% for *Ca. Bgiplasma* family(**Supplementary Table 5**). We found *Ca. Bgiplasma* are gene content dominated by replication, recombination and repair, posttranslational modification, protein turnover, chaperones, and inorganic ion transport and metabolism, which are reported active up-regulated in *Deinococcus* during gamma-irradiation^40^ (Supplementary Fig. 2b).

### Distribution of species and strains

The new samples from this study differed in oral microbiome composition compared to published samples across geography/ethnicity (Fig. 5a,b). Both the 4D-SZ and Yunnan samples abundantly contained many uSGBs (of the top 10 abundance species in cohort) such as *Neisseria* spp., *Porphyromonas* spp. and kSGBs such as *Haemophilus parainfluenzae* and *Veillonella denticarios*, which were rare in the other cohorts (Fig. 5b). Pregnant samples from the U.S. contained *Fannyhessea vaginae* (the vaginal pathogen previously known as *Atopobium vaginae*^41^), *Urinacoccus*, etc. that were of much lower abundance in other cohorts (Fig. 5b). Samples from Fiji, although not well mapped (Fig. 2a), showed high levels of a few SGBs that were also seen in the RA study from Beijing, China, including an SGB from *Saccharibacteria* (TM7) (Fig. 5b).

**Figure 5.**
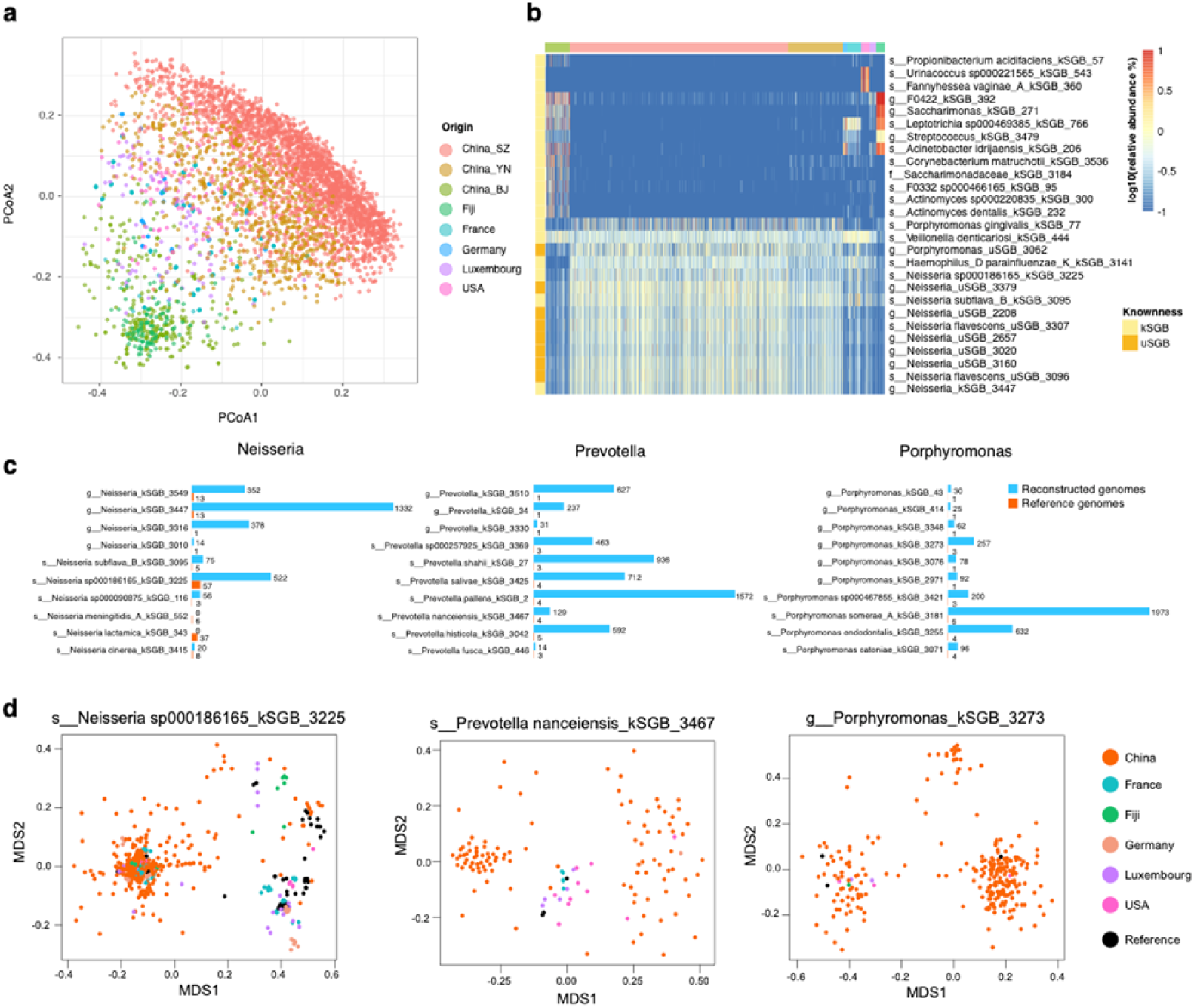
Geographical distribution of oral SGBs and strains. **a**, Principal coordinate analysis plot based on Bray-Curtis distances of oral SGB relative abundance profile highlights distinct microbial communities among different origin populations. **b**, The relative abundance of top 10 most abundance SGBs from each origin populations. A large set of reconstructed uSGBS are widely high abundance distribution in our cohort (Shenzhen and Yunnan) and lack of several highly abundant kSGBs in other population. Species are order by hclust with complete linkage and euclidean distance. **c**, Our reconstructed MAGs largely extend the size (genome numbers) of the top 10 most abundance species from common oral genus with few reference genomes. **d**, Multidimensional scaling on average nucleotide identity between MAG and reference genomes in species showed strain variety and that constructed MAGs dominated the sub species. Only HQ MAGs and reference genomes are showed.

At the strain level, the new samples from the current study greatly expanded the genome collection for common taxa such as *Neisseria* spp., *Porphyromonas* spp., and *Prevotella* spp. (Fig. 5c,d). The numbers of publicly available reference genome for the top ten most abundant species in the genera *Porphyromonas* and *Prevotella* were less than 10, and less than 100 for the genus *Neisseria*. Here we obtained more than 1000 genomes for a few of the species, and increased the diversity in all the species in these genera (Fig. 5c). Most of the species with a large number of genomes showed strain-level variations (subspecies). The *Prevotella nanceiensis* kSGB, for example, included 3 reference genomes that were similar to a few genomes from developed countries, while our samples contributed two large clusters that were more distantly related (Fig. 5d).

### New disease markers according to the oral genomes

To illustrate the utility of our genome collection in metagenomic studies including MWAS, we reanalyzed dental and salivary microbiome data from RA patients and controls^6^. For better confidence in the markers regardless of cohort, we only analyzed SGBs containing >10 genomes. Similar to the original study, oral markers selected by a 5x 10-fold cross-validated gradient boosting algorithm include a number of Gram-negative bacteria e.g. *Haemophilus* spp., *Aggregatibacter* spp. enriched in dental samples from healthy volunteers, while only a *Pseudomonas* SGB and a *Enterococcus* SGB were selected for RA samples (Fig. 6a). Interestingly, the two new RA dental markers appeared more abundant in control saliva samples. The strongest marker from healthy saliva remained *Lactococcus lactis*^5^, and *Lactobacillus paracasei, Streptococcus infantarius, w*ere identified, reminiscent of beneficial effects of *L. casei* gavage in rat model of RA^42, 43^. The assembled genomes allowed matching of different species in the *Veillonella* genus as RA saliva markers. Moreover, *Pauljensenia* spp., a genus recently renamed from *Actinomyces*^44^, was identified as highly predictive of RA. As *Actinomyces* are the basis for dental attachment of oral bacteria^45^, potential contribution of *Pauljensenia* spp. to periodontitis in RA patients remains to be explored; the dental microbiome was obviously deranged, consistent with epidemiology^5^.

**Figure 6.**
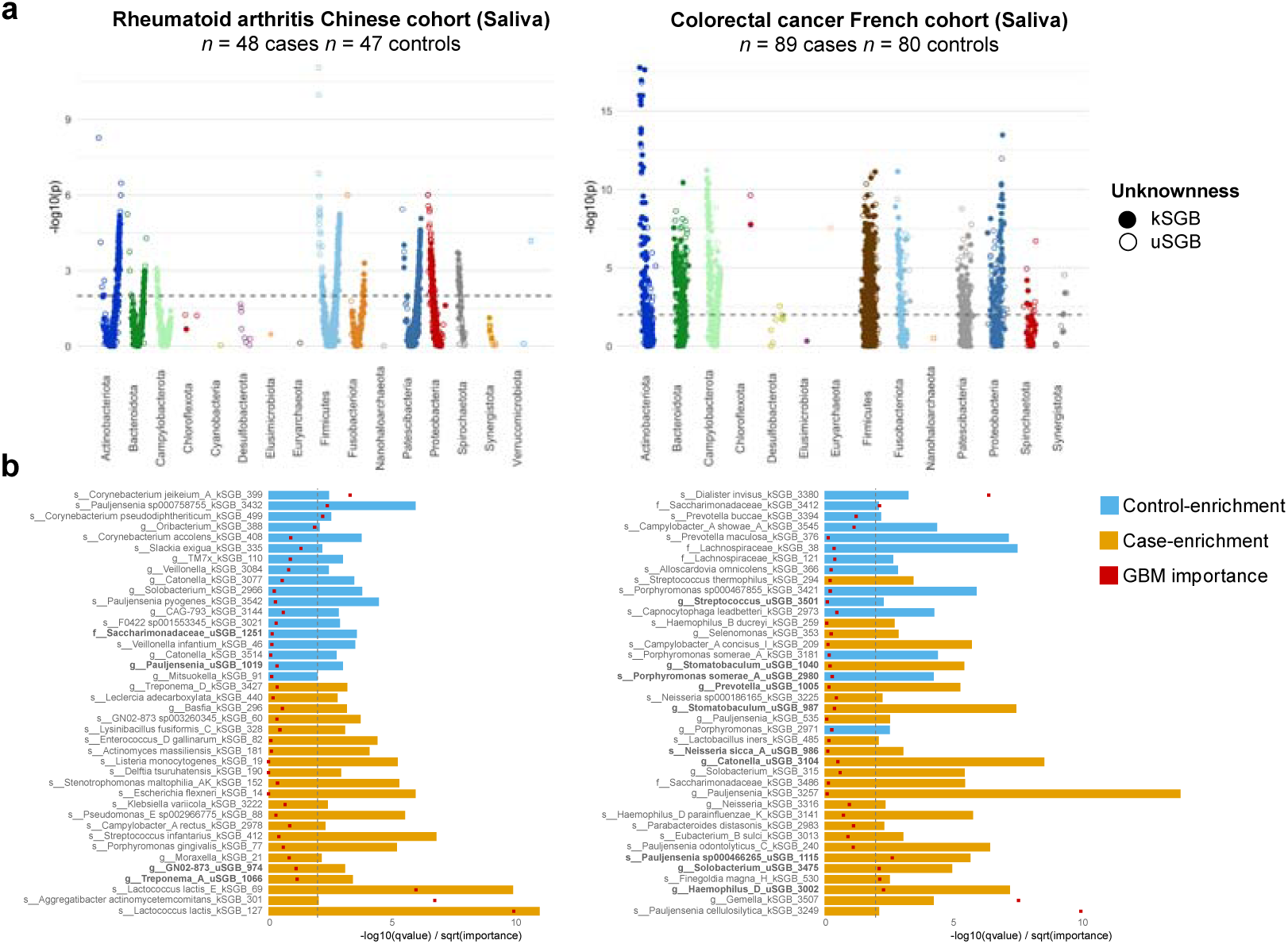
Disease markers according to the oral genomes. **a**, The Manhattan plot shows metagenomic wise association of oral SGBs for RA and CRC studies. The species are ordered according to their phylogeny (bottom) and the association direction (positive or negative). Each point is one SGB and point height indicates the FDR value correction for multiple hypothesis tests from a generalized linear model (GLM) test between diseased and healthy species abundance after adjusting Age, Gender, BMI. The dotted line indicates a false discovery rate (FDR) of 1%. **b**, SGBs association with RA and CRC. We select 40 large and most importance oral SGBs (>10 genomes) for disease prediction using Gradient Boosting Machine (GBM). The species are order according to their partial spearman correlation adjusted age, gender, BMI and GBM importance. The bar length indicated the FDR value between groups as described above. The dotted line indicates a false discovery rate (FDR) of 1%. The red square in bar is the sqrt GBM importance. uSGBs are highlight in bold label text.

A set of saliva samples from colorectal cancer and controls from France are also available^46^. Here, we found *Pauljensenia* spp., to be control-enriched, along with *Acinetobacter radioresistens*, *Lachnoanaerobaculum* sp., *Catonella* sp., etc (Fig. 6b). *Streptoccocus thermophilus*, a species previously found to be enriched in fecal samples from control or adenoma compared to CRC patients^47^ was also identified in control saliva. The markers enriched in CRC oral samples are more unexpected. Besides *Porphyromonas* spp., *Prevotella maculosa*, we found a *Lachnospiraceae* SGB (potentially TMA-producing and consistent with gut results^10, 48–50^), *Capnocytophaga leadbetteri, Cardiobacterium hominis, etc.* (Fig. 6b). Thus, the substantially expanded collection of oral microbial genomes enabled discovery of new disease markers and genomic representation of previously reported markers, facilitating the shift from fecal to oral microbiome-based diagnosis and therapeutics.

### Potential functions in drug metabolism and small molecule synthesis

Many human target drugs are reported to be metabolited to its inactive form by gut human microbiome^51, 52^ or impact the gut bacteria^51–53^. Gut bacteria genes that metabolite 41 human targeted drugs, 6 non-traditional antibacterial therapeutic and key enzymes experimented validated for 12 human diseases were mapped to our oral SGB genomic contents (**Supplementary Tables 6**). We show that many oral communities share homologous to these gut bacteria encoding enzymes, suggesting the oral microbiome may also play an importance role in medical therapy and disease development (Fig. 7a). More specificly, there are total 2696 SGBs containβ-glucuronidase enzyme that can metabolite anti-cancer drug Gemcitabine (2’, 2’-difluorodeoxycytidine) into its inactive form^52^. 456 SGBs have agmatine gene for anti T2D drug Metformin and 225 SGBs have tyrosine decarboxylase (TyrDC) for anti-parkinson drug L-dopa^51^. There are also 1733 oral SGBs have genes producing small molecule taurine and 5-aminovalerate which are potential drugs for autism spectrum disorder (ASD)^51^. Unexpected few SGBs contain CutC/CutD genes which are key enzyme for TMA, a metabolite with high cardiovascular event risk^51^.

**Figure 7.**
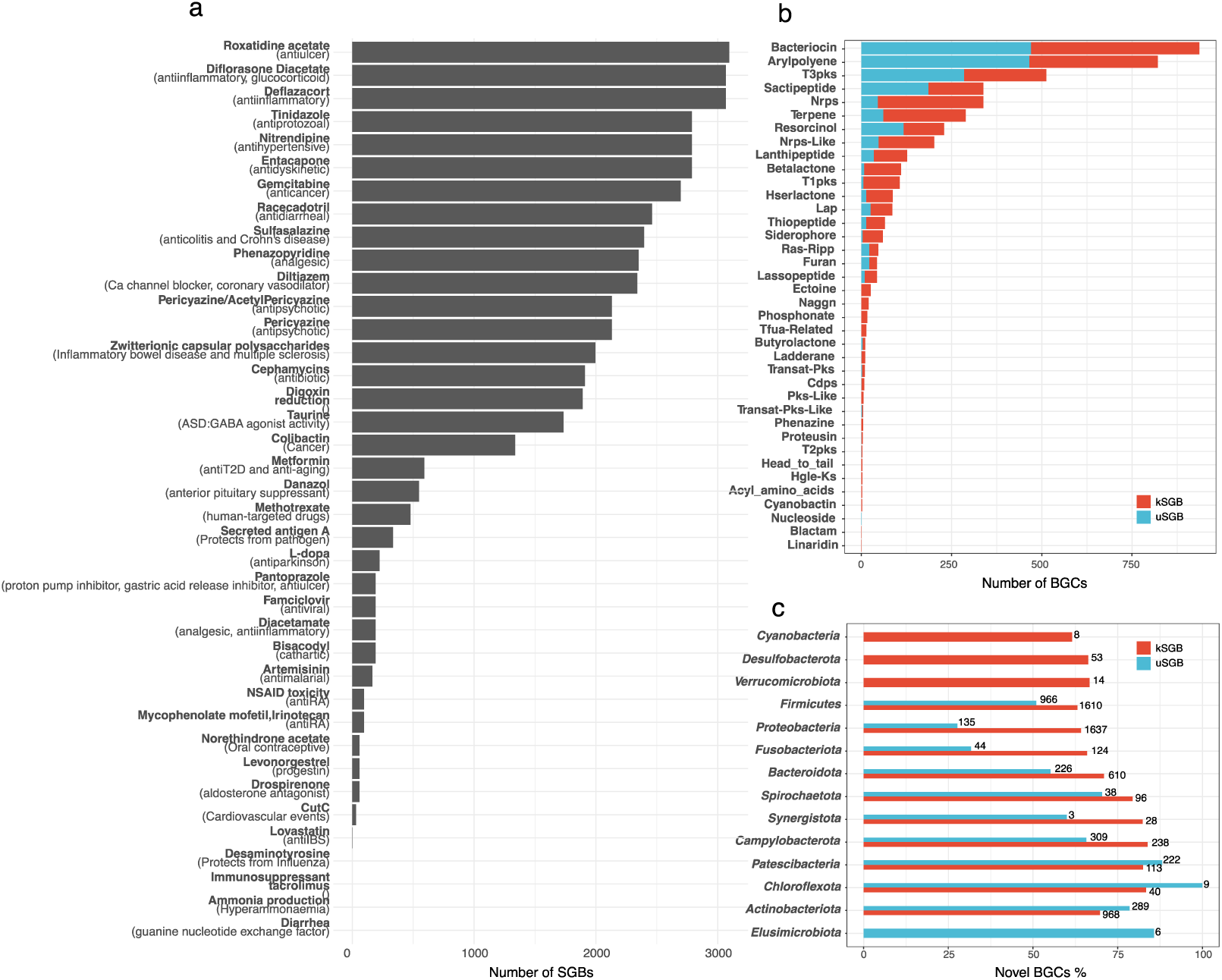
A comprehensive mapping of the function repertoire of the human oral microbiome. **a**, Number of oral SGBs who share homologous to gut microbiome enzyme coding drug metabolite or healthy related function. Details see Supplementary Table 6. **b**, Summary of predicted BGCs in oral microbiome. Numbers of BGC of 38 different types detected in the 2711 oral bacterial genomes were grouped by kSGB and uSGB. **c**, Fraction of novel BGCs across phylum levels. The numbers of bar show the novel BGC count while bar length represent kSGBs/uSGBs ratio on the phylum levels.

The inferration of secondary metabolites biosynthetics gene clusters (BGCs) was made by applying antiSMASH^54^ pipeline. The total 12399 BGCs (7804 unknown, 4595 known) have been detected from 91.46%(1167) kSGBs and 66.75%(1544) uSGBs, and the BGCs coding for bacteriocin, arylpolyene, type III **PKS** (polyketide synthase) has appeared more than 500 times on the oral bacterial community(Fig. 7b**, Supplementary Table 7**). For each specie’s genome, the size percentage (mean: 2.512%) of BGCs was calculated based on the each BGC’s location on the each genome. The vast majority of the genome has a BGC range of less than 10% compared to the total genome (Supplementary Figure 5), included *Firmicutes*, *Patescibacteria*, *Actinobacteriota*, *Proteobacteria*, etc. Notably, there represented 67% novel BGCs (2743 known, 5557 unknown) in the kSGBs and 54% novel BGCs (1852 known, 2247 unknown) in the uSGBs. At the phylum level, *Elusimicrobiota*, *Actinobacteriota*, *Chloroflexota*, *Patescibacteria* contains a higher proportion of novel clusters (Fig. 7c). These unknown BGCSs demonstrate the enormous potential of oral microbes for the synthesis of natural metabolites for drug development and disease treatment.

## DISCUSSION

In summary, we provide the largest set of oral metagenomic shotgun data, assemble tens of thousands of draft genomes for the human oral microbiome, including 2,313 new species as well as many new strains of known species. The results illustrate that culturomics have not even exhausted the microbial complexity in the more accessible body sites, and that metagenomic data for large cohorts of non-fecal samples have great potential. A number of taxa with compact genomes were identified in this study, such as CPR and Mollicutes. Mollicutes such as *Mycoplasma* and *Ureaplasma* are well known in the female reproductive tract^22^. Much remains to be elucidated for the metabolic requirement of small bacteria in the oral microbiome. Oral bacteria also contributed to discovery of new CRISPR-Cas systems^55^. Species with thousands of metagenomic and isolated genomes would be amenable to microbial GWAS^56^ (microbial genome-wide association studies) to discover virulence factors, drug resistance and more commensal functions, which has so far only be possible for pathogens.

### Accession codes

All the data are available at China National Genebank (CNGB), Shenzhen under the accession CNP0000687. https://db.cngb.org/microbiome/

## AUTHOR CONTRIBUTIONS

J.W. initiated the overall health project, H.J. decided to include oral samples, Z.J., J.Z., L.T. worked out the metagenomic assembly approach, with Hadoop support from X.L. M.H., Z.L., C.L. contributed Yunnan samples. P.C., K.C., X.W., Y.L. contributed 4D-SZ samples, and L.S., X.T. managed the samples and data. J.Z., L.T., Z.J., H.J. interpreted the data, prepared the display items and wrote the manuscript. All authors contributed to finalizing the manuscript.

## COMPETING FINANCIAL INTERESTS

The authors declare no competing financial interest.

## ONLINE METHODS

### The newly cohort and published datasets used in this study

The 2675(2284 saliva and 391 tongue) oral metagenomics samples from Chinese 4D-shenzhen corhorts(**Supplementary Tables 1 sheet 4**) and the 671 salivary samples from six cities and villages in Yunnan province were collected in this study(**Supplementary Tables 1 sheet 5**), and total 706 public oral metagenomics datasets^6, 23–26^ were downloaded from NCBI SRA databases with accession codes SRP029441, ERP006678, SRP133047, ERP110622 and SRP07256, encompassing five different studies (**Supplementary Table 1 sheet 3**) have been reported previously.

### Sample collection, DNA extraction, sequencing and quality control

The 2955 salivary samples and 391 tongue samples from Shenzhen were self-collected by volunteers, using a kit containing a room temperature stabilizing reagent to preserve the metagenome^57^. DNA extraction of the stored samples within the next few months was performed using the MagPure Stool DNA KF Kit B (MD5115, Magen) from 1mL of each sample. Metagenomic sequencing was done on the BGISEQ-500 platform^58^ (100bp of paired-end reads for all samples and four libraries were constructed for each lane) and generated 101.4 billions pairs of raw reads. The 671 salivary samples from Yunnan province were self-collected using commercial kits (Cat. 401103, Zeesan, China). Collected samples were temporarily stored in −80℃ freezers and then transported to CNGB, Shenzhen with dry ice via commercial logistics (SF Express Inc.). DNA was extracted in the same way as above. Sequencing was performed on the BGISEQ-500 machines and generated 26.5 billions single-end 100 bp length reads. The raw read length for each end was 100bp. After using the quality control module of metapi pipeline followed by reads filtering and trimming with strict filtration standards(not less than mean quality phred score 20 and not shorter than 51bp read length) using fastp v0.19.4^59^, host sequences contamination removing using Bowtie2 v2.3.5^60^ (hg38 index) and seqtk^61^ v1.3, we totally got 54.9 billions high-quality PE reads and 7.1 billions high-quality SE reads.

### Metagenomic De novo assembly, binning and checkm

The high-quality PE and SE reads was individually assembled using assembly module of metapi pipeline with different max kmer cutoff by different max read length of each samples applying SPAdes v3.13.0^28^ (PE reads with option --meta^27^). All configuration can see on https://github.com/ohmeta/metapi/blob/dev/metapi/config.yaml. After we got draft genomes on contig level of each samples, the reads was mapped back to each assemblies using BWA-MEM v0.7.17^62^ with default parameters and calculate the contig depth by jgi_summarize_bam_contig_depths^21^, then using MetaBAT2 v2.12.1^21^ to do metagenomic binning individually for each samples. Finally we got totally 163,718 bins. After MAGs quality assignment by CheckM v1.0.12^30^ lineages workflow, 15,013 high-quality (completeness > 90% and contamination < 5%, HQ) bins and 41,200 medium-quality(completeness > 50% and contamination < 10%, MQ) bins(**Supplementary Table 2**) have been generated based on MIMAG standard^29^. The 16S rRNA sequences in the MAGs were searched by Barrnap v0.9^63^ with parameters “--reject 0.01 --evalue 1e-3” and tRNA sequences in the MAGs were searched by tRNAscan-SE 2.0.3^64^ with the default parameters.

### Public database used

The public bacteria and archaea genomes database used in this study include(**Supplementary Table 1 sheet 6**):

a. The NCBI Refseq bacteria and archaea databases (ftp://ftp.ncbi.nlm.nih.gov/genomes/refseq/, accessed in June 2019) contain 155854 microbial genomes.
b. The eHOMD^31^ database (http://www.homd.org/ftp/HOMD_prokka_genomes) contain 1526 microbial genomes come from human oral environment.
c. The IGGdb^15^ (https://github.com/snayfach/IGGdb) contain 23790 microbial genomes come from human gut environment.
d. The hSGBRep^16^ database contain 4930 representative microbial genomes (http://segatalab.cibio.unitn.it/data/Pasolli_et_al.html) come from human body site include gut, oral, skin, genital tract.
e. The BPUMGs^14^ databases (ftp://ftp.ebi.ac.uk/pub/databases/metagenomics/umgs_analyses/) contain 1952 microbial genomes come from human gut.
f. The CGR^33^ database accession code <u>PRJNA482748</u> contain 1520 microbial genomes come from human gut bacterial culture collection.
g. The HBC^32^ database (ftp://ftp.ebi.ac.uk/pub/databases/metagenomics/hgg_mags.tar.gz) contain 737 microbial genomes come from human gut bacterial culture collection.

### Clustering metagenomic genomes into species-level genome bins

The 56,213 reconstructed genomes and 190,309 reference genomes were grouped into species-level genome bins(SGBs) by a two-step clustering strategy as reported previously^16^ with a slight modification. In the first step, all-versus-all genetic distance matrix between the 246,522 genomes was carried out using Mash version 2.0^65^ (“-k 21 -s 1e4” for sketching). Then, hierarchical clustering with average linkage and 0.05 genetic distance cutoff on the distance matrix by fastcluster^66^ was resulted to 33008 clusters. Because the Mash will underestimate the distance between the incomplete genomes^67^ and split same-species genomes into multiple SGBs, we performed clustering base on average nucleotide identity(ANI) in the second step. First, We divide the SGB into known SGB(kSGB) and unknown SGB(uSGB) according to with or without reference genomes. Then, a representative genome was selected for each SGBs. For the kSGB, the genome which has the largest genome size was selected. For the uSGB, all MAGs were rank by completeness(in decreasing order), contamination(increasing), coverage(decreasing), strain heterogeneity(increasing), N50(decreasing). And representative genome was selected as the one minimizing the sum of the five ranks. We recalculated the more precise genetic distance using pyani v0.2.9^68^(option ‘-m ANIb) for the pairs of representative genomes with mash distances less than 0.95 and only left ANI with genome coverage above 0.3. Following hierarchical clustering with complete linkage based on >95% ANI score, 12,911 representative genome which mash distances less than 0.95 were merged to 11,427 new clusters. Finally, we obtained 31,525 SGBs by two-step clustering strategy. In this dataset, only 3,589 SGBs included eHMOD genomes and oral metagenomes MAGs were named Oral SGBs and can be further divided into 2,313 uSGBs and 1,276 kSGBs. The top three contributions of uSGBs are 4D_SZ (1441 uSGBs), ZhangX_2015 (445 uSGBs) and Yunnan (334 uSGBs). The other 27,936 SGBs are non-oral SGBs (Figure 1b).

### Reconstruction of the human-oral microbiome phylogenetic structure

The phylogenetic trees of 3589 representative genomes of SGBs (Figure 3C) and 76 genomes of Acholeplasmataceae Order were both built using the 400 PhyloPhlAn markers with the parameters “--diversity high --fast --min_num_markers 80” by the PhyloPhlAn2^69^. As input data for PhyloPhlAn2, proteome were predict using Prodigal v2.6.3^70^ with default parameters. Following tools with their set of parameters were used in the configuration files:

Diamond v0.9.22.123^71^ with parameters: “blastp --quiet --threads 1 --outfmt 6 --more-sensitive --id 50 --max-hsps 35 -k 0”;

Mafft v7.407^72^ with the “--anysymbol” option;

Trimal v1.4.rev15^73^ with the “-gappyout” option;

Iqtree v1.6.12^74^ with parameters: “-quiet -nt AUTO -m LG”.

The phylogenetic trees in figure 3c was generated using GraPhlAn v1.1.3^75^ and the phylogenetic trees in figure 4a, figure s2 were generated using FigTree v1.4.4 (https://github.com/rambaut/figtree/releases).

### SGBs taxonomic and function analyses

The taxonomic classification of 3589 representative genomes of SGBs was assigned using GTBD-Tk v0.3.2^70, 76–79^ (https://github.com/Ecogenomics/GTDBTk) classify workflow with external GTDB database release 89.0(https://data.ace.uq.edu.au/public/gtdbtk/release_89/89.0/). Although some kSGBs already have taxonomy label, we still using GTDB-Tk to classify them because GTDB-Tk has its own taxonomy classification system that is different from the NCBI taxonomy database. Then above the genus level, we manually removed the classification tag with a single letter suffix (**Supplementary Table 3**). Those suffixes used to indicate that taxon needed to be subdivided based on the current GTDB reference tree. We used EggNOG mapper v1.0.3^39^ to do genome-wide functional annotation through orthology assignment on 3589 SGBs(**Supplementary table 3**) and 29 MAGs in Candidatus bgiplasma (Supplementary Figure 2b). The secondary metabolite biosynthesis gene clusters(BGCs) of 3587 oral bacterial genomes was identified respectively by using antiSMASH v5.0.0^54^ with options --fullhmmer --cf-create-clusters --smcog-trees --cb-knownclusters --asf --pfam2go. Then we use the bgctk(https://github.com/ohmeta/bgctk) to parse and merge BGCs’s results from all json file which was generated by anstiSMASH workflow.

### Mapping rate compared between different oral related genomes database

The mapping rates of oral metagenomics reads align to three different oral related genomes databases(eHOMD, hSGB_Rep, oralSGB_Rep) were compared based on the statistics summary of Bowtie2’s results(**Supplementary Table 4**). First we randomly selected 100 oral metagenomes samples from each of 4D_SZ and Yunnan cohorts. With all 808 public samples and 81 additional verify samples which not used to assembly (**Supplementary table 1 sheet 2**), the total 1089 oral metagenomes samples were mapped to these databases respectively using Bowtie2 v2.3.5 with default parameters. The barplot of mapping rate was generated using R package ggplot2 3.1.1^80^ faced with different databases and different country.

### Metapi for oral SGB metagenomic profiling

The quantification of species relative abundance of oral metagenomic samples was performed with the taxonomic profiling module of metapi pipeline: i) build the oral representative SGBs’ index by Bowite2; ii) align the high-quality reads of each sample to the oral genome index using Bowtie2 with parameters: “--end-to-end --very-sensitive --seed 0 --time -k 2 --no-unal --no-discordant -X 1200”; iii) The normalized contigs depths were obtained by using jgi_summarize_bam_contig_depths; vi) base on the correspondence of contigs and genome, the normalized contig depth were converted to the relative abundance of each SGB for each samples. Finally we merged all representative SGBs relative abundance to generate a taxonomic profile.

### PCOA, heatmap and oral type for metapi profile

Principal Coordinates Analysis (Pcoa) of metapi profile was done used dudi.pco function in ade4^81^ R package based on bray distance from vegan2.5.2^82^ R package. The mean top 10 most abundance SGBs from every study were merged (total 27 SGBs) to visual in pheatmap^83^ R package.

### Pangenome, phylogenetic analysis of kSGB and uSGBs

From the taxonomic profiling results of 4820 oral metagenomic samples, the most prevalent eight genus was selected based the rank of average relative abundance(decreasing), occurrence frequency(decreasing), oral genome number / SGBs size(decreasing), include *Prevotella*, *Neisseria*, *Streptococcus*, *Veillonella*, *Porphyromonas*, *Fusobacterium*, *Pauljensenia*, *Haemophilus*. Then we choose ten most prevalent species for each genus to do pangenome analysis. First each species has a representative genome correspond to each SGBs, so we use prokka v1.13.7^84^ to do genome annotation for all genomes of each SGBs. Then the annotated genomes were used to construct pangenome database for each SGBs via panphlan_pangenome_generation.py (a script come from PanPhlAn v1.2^85^). Finally the gene-family presence / absence profile matrix was transformed to a zero/one matrix for reference genomes and reconstructed genomes of each SGBs to do rarefaction analysis. Accumulation curves (Supplementary Figure 3) based on the number of core gene of each SGBs were bootstrapped ten times at each sampling interval. The observation of intra-SGB phylogenetic structure of *Neisseria* kSGB 3225, Prevotella kSGB 3467 and *Porphyromonas* kSGB 3273 was performed by the nonmetric multidimensional scaling analysis using the metaMDS function of R package vegan v2.5.2.

### Disease markers according to the oral genomes

The metagenomics wise association between 3,589 metapi species profiles (SGB) and disease for previously published CRC and RA studies was done using generalized linear model (GLM) with adjust for potential confounders such as gender, age, BMI (Table S1). BMI is only available for RA. Species relative abundances was asin-sqrt transformed as described before^86^. Non-oral SGBs were excluded. Corrected for multiple hypothesis tests was done using FDR. We predicted disease status using gradient boosting model (GBM) in caret^87^ R package, such that 80% of the samples were randomly sampled for each estimator. The depth of the tree at each estimator was not limited, but leaves were restricted to have at least 30 instances. We used 4000 estimators with a learning rate of 0.002. All the FDR <1% oral marker SGBs are included in the model as predictors. To avoid overfitting, 5 repeat ten folds cross validation ROC was used to measure the model performance. VarImp function was used to extract the GBM importance.

## ACKNOWLEDGEMENTS

This research was supported by the Shenzhen Municipal Government of China (), Guangdong (). We gratefully acknowledge colleagues at BGI-Shenzhen and China National Genebank (CNGB), Shenzhen for sample collection, DNA extraction, library construction, sequencing, and discussions.

## Supplementary Figures

**Supplementary Figure 1.**
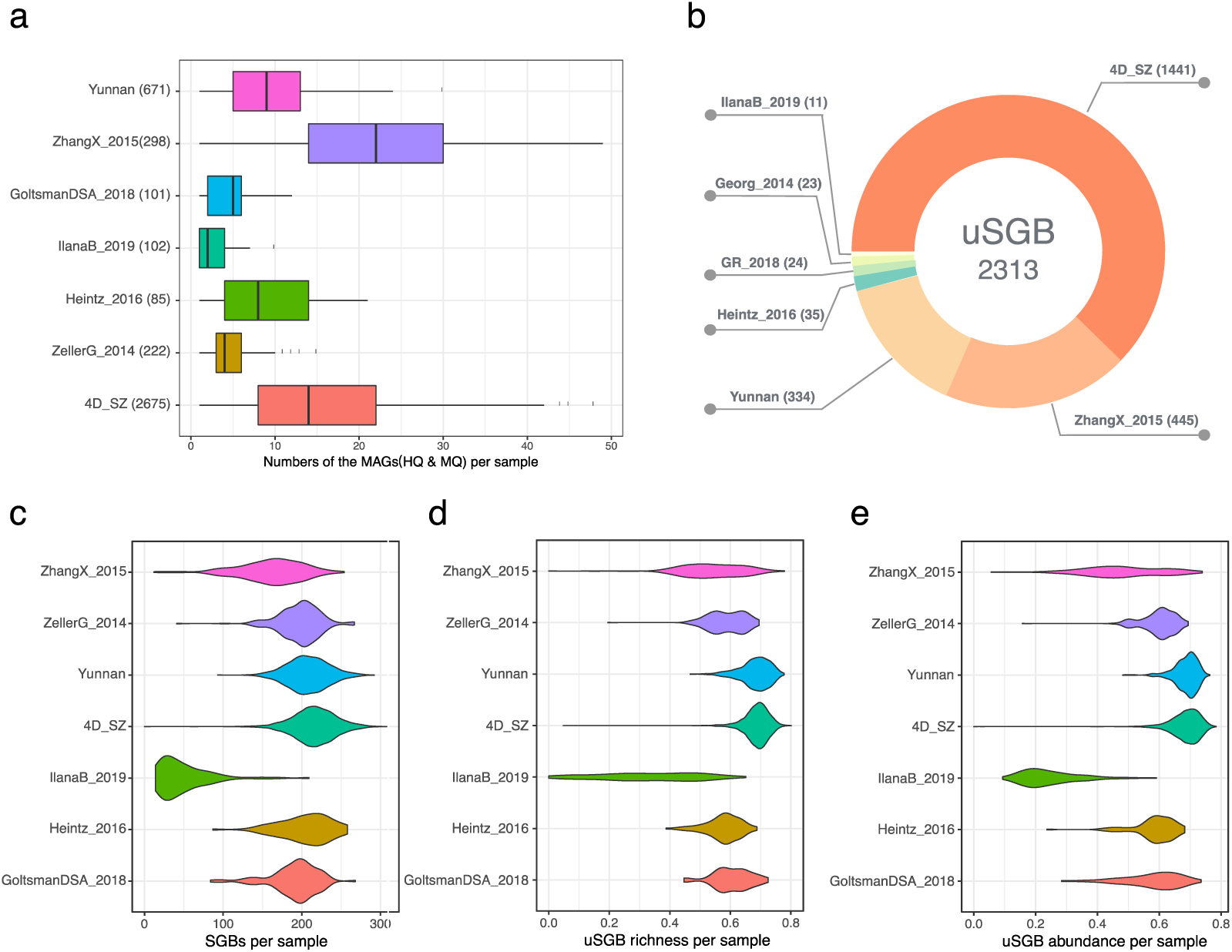
Summary of assembly quality, SGBs distribution across 7 studies. **a**, Numbers of medium quality and high quality MAGs in samples from 7 studies. **b**, 3,589 uSGBs origin distribution across studies. **c**, The number of all SGBs (>0.001 abundance) for each samples. **d**. uSGB richness (number of uSGB/number of all SGB) for each sample. e. Sum of all uSGB abundance for each samples.

**Supplementary Figure 2.**
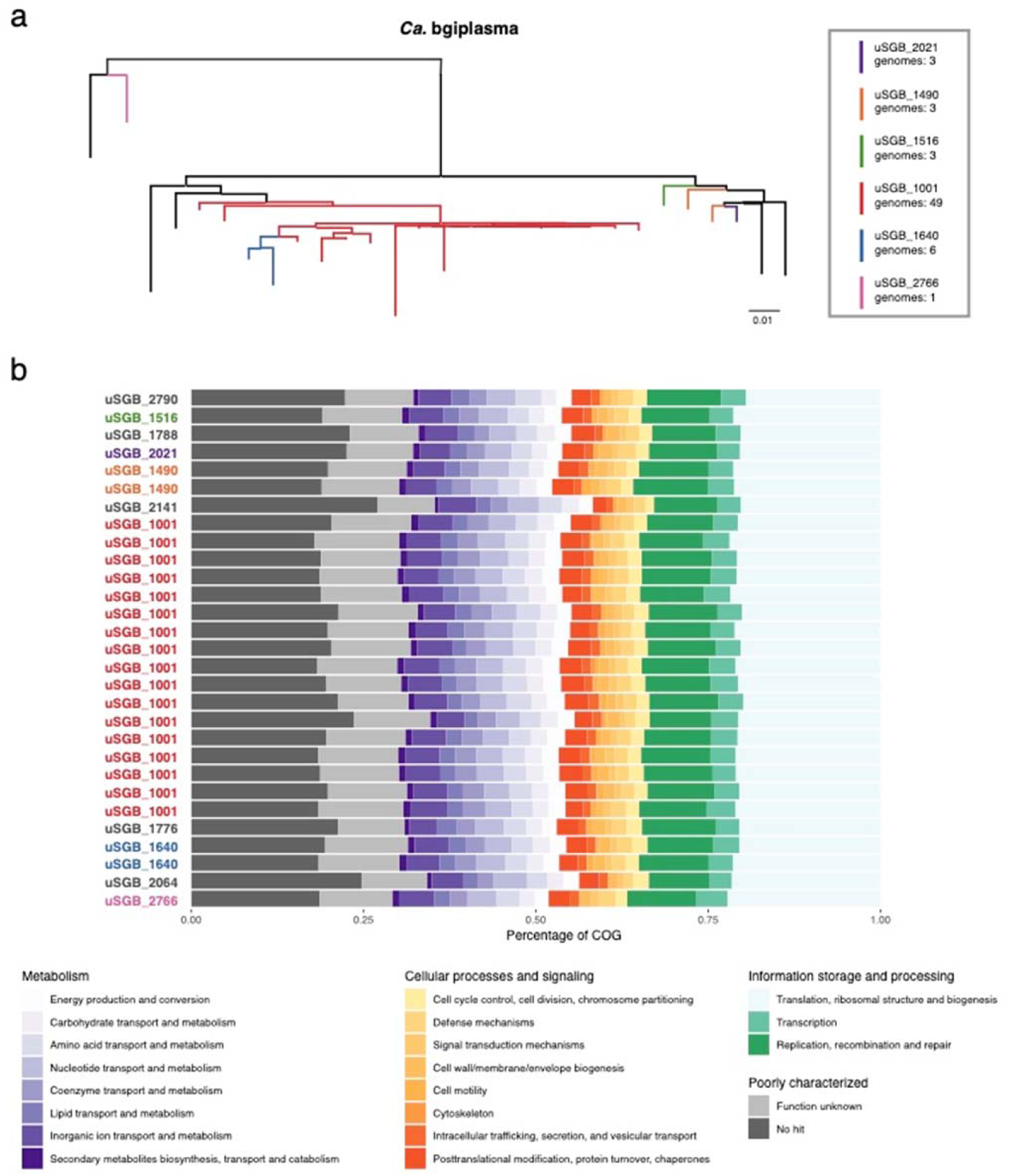
Phylogenetic tree and COG functional annotation from all MAGs in the new candidatus family. **a**, A phylogenetic tree from *Ca*. bgiplasma (Fig. 4a) are displayed detailed here. The high quality MAGs belong the same species are colored the same color and medium quality MAGs are left black. **b**, *Ca.* bgiplasma function genome annotated by EggNOG mapper. The main function category of COG are displayed as the percentage of genes annotated to that category.

**Supplementary Figure 3.**
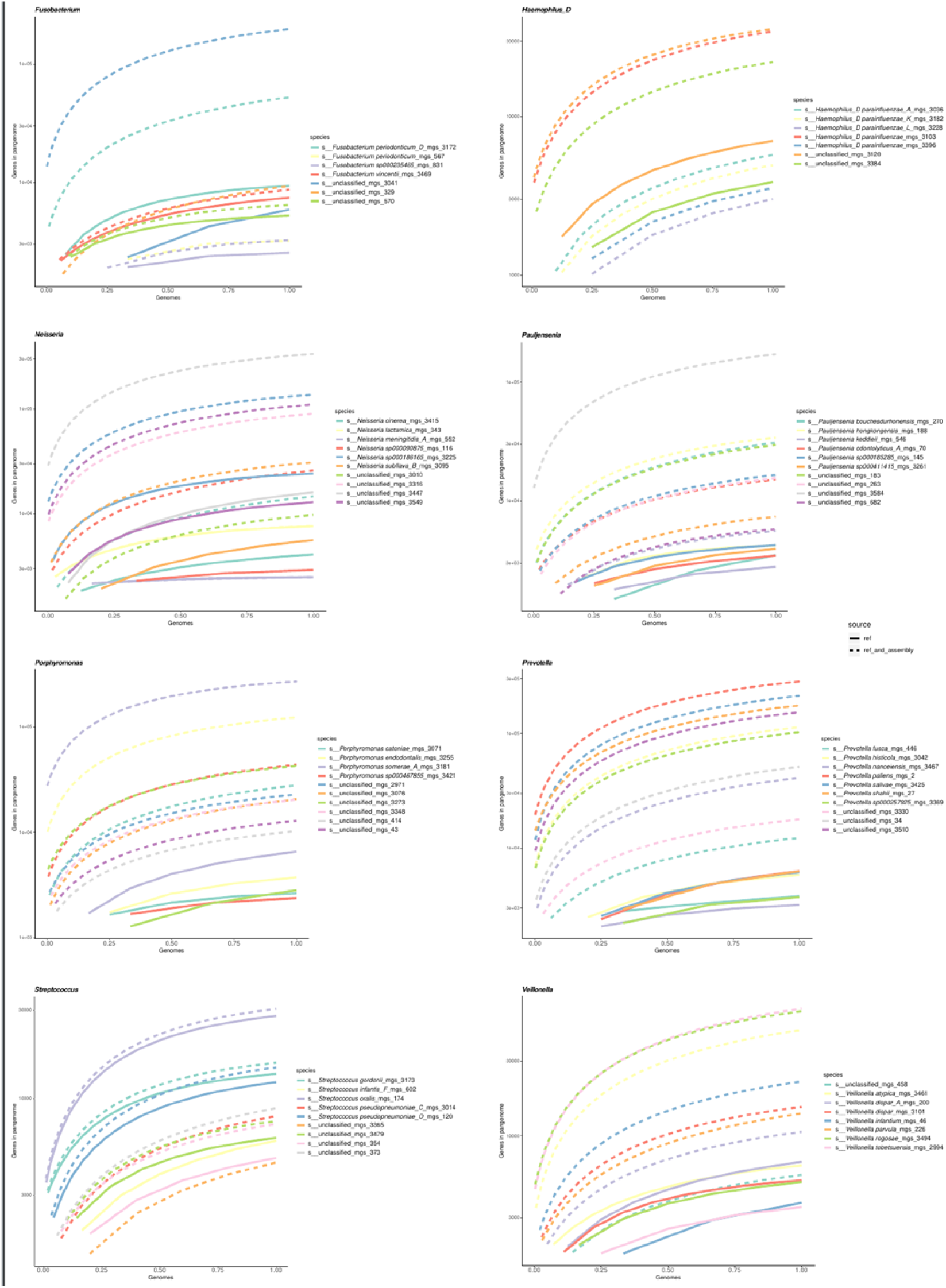
The pangenome genetic diversity. The gene number rarefaction curve indicated that the genetic diversity have been increased through more genomes metagenomically assembled, included 71 species genomes come from eight most prevalent genus.

**Supplementary Figure 4.**
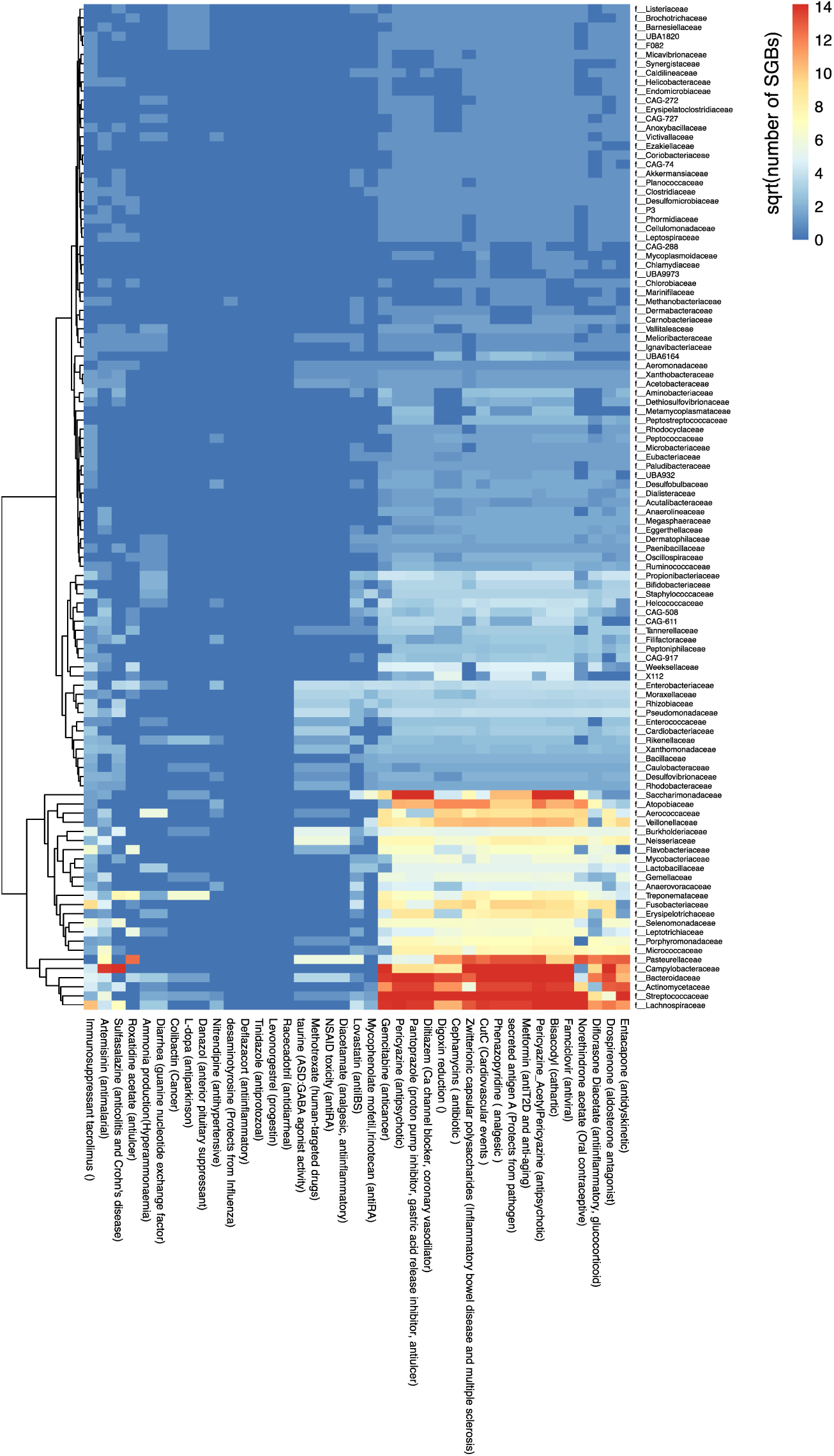
Heatmap of number of SGBs at family level who share homologous to gut microbiome enzyme coding drug metabolite or healthy related function. Cell is sqrt number of SGBs. Hclust with canbera distance.

**Supplementary Figure 5.**
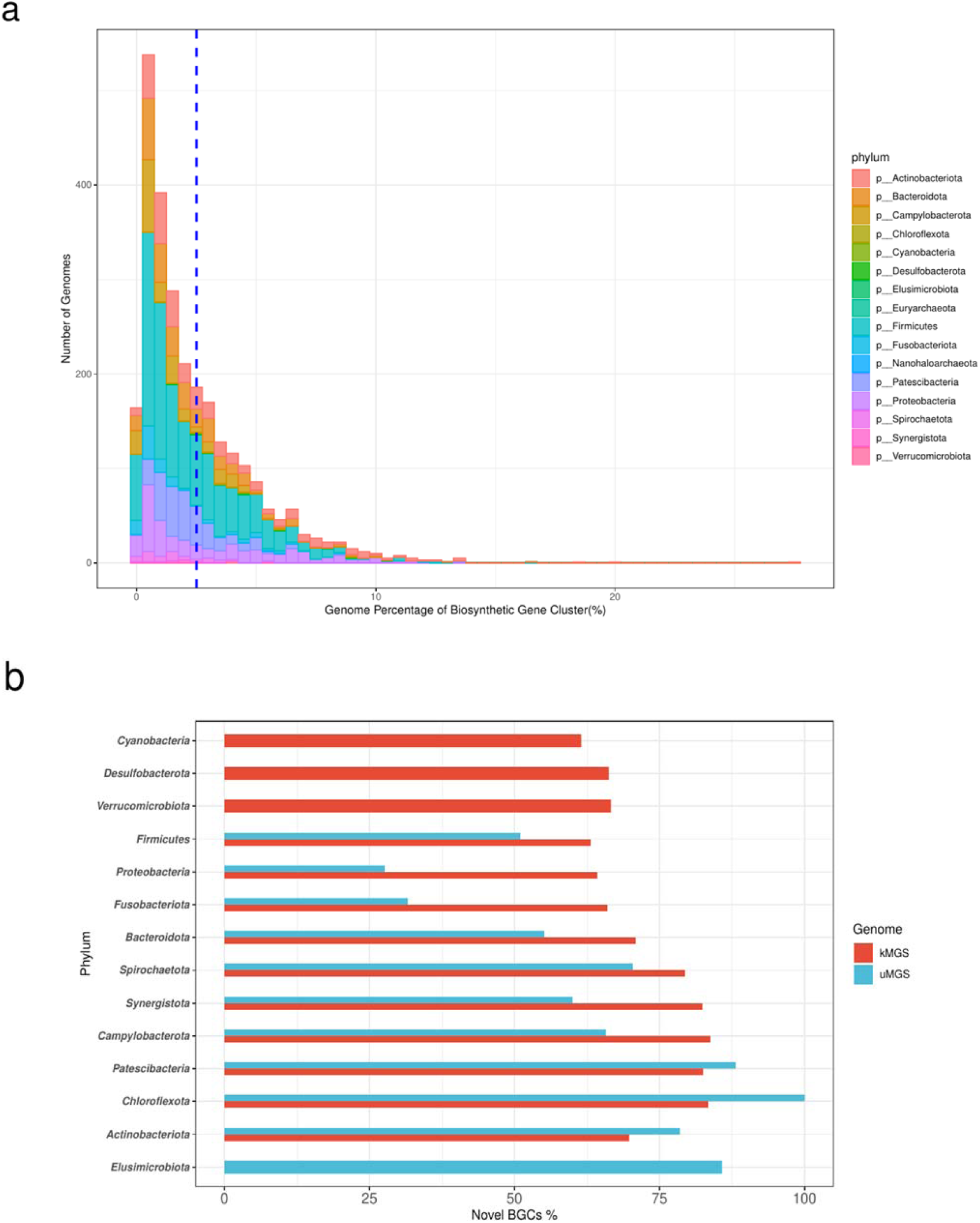
The detection of BGCs on the human oral microbiome. **a**, The distribution of genome percentage of biosynthetic gene cluster. From oral microbiome included 2713 genomes form 16 different phylum. The x coordinate corresponds to the proportion of BGC size, and the y coordinate corresponds to the number of genomes. The blue vertical line indicates the average proportion of BGC size: 2.512%. **b**, Novel BGCs proportion on phylum level of oral microbiome. The x coordinate corresponds to the proportion of the number of the novel BGCs, the y coordinate corresponds to the 14 different phylum groupped by uSGB and kSGB.

## References

1. Cabreiro, F. & Gems, D. Worms need microbes too: microbiota, health and aging in Caenorhabditis elegans. EMBO Mol Med 5, 1300–1310 (2013).

2. Zhernakova, A. et al. Population-based metagenomics analysis reveals markers for gut microbiome composition and diversity. Science 352, 565–569 (2016).

3. Jie, Z. et al. A multi-omic cohort as a reference point for promoting a healthy gut microbiome. To be Submitt. soon (2019).

4. Methé, B.A. et al. A framework for human microbiome research. Nature 486, 215–221 (2012).

5. Lloyd-Price, J. et al. Strains, functions and dynamics in the expanded Human Microbiome Project. Nature 550, 61–66 (2017).

6. Zhang, X. et al. The oral and gut microbiomes are perturbed in rheumatoid arthritis and partly normalized after treatment. Nat Med 21, 895–905 (2015).

7. Qin, N. et al. Alterations of the human gut microbiome in liver cirrhosis. Nature 513, 59–64 (2014).

8. Wang, J. & Jia, H. Metagenome-wide association studies: fine-mining the microbiome. Nat Rev Microbiol 14, 508–522 (2016).

9. Atarashi, K. et al. Ectopic colonization of oral bacteria in the intestine drives TH1 cell induction and inflammation. Science 358, 359–365 (2017).

10. Jie, Z. et al. The gut microbiome in atherosclerotic cardiovascular disease. Nat Commun 8, 845 (2017).

11. Yu, J. et al. Metagenomic analysis of faecal microbiome as a tool towards targeted non-invasive biomarkers for colorectal cancer. Gut 66, 70–78 (2017).

12. Liu, X. et al. M-GWAS for the Gut Microbiome in Chinese Adults Illuminates on Complex Diseases. Cell (2019).

13. Chen, T. et al. The Human Oral Microbiome Database: a web accessible resource for investigating oral microbe taxonomic and genomic information. Database (Oxford) 2010, baq013 (2010).

14. Almeida, A. et al. A new genomic blueprint of the human gut microbiota. Nature 568, 499–504 (2019).

15. Nayfach, S., Shi, Z.J., Seshadri, R., Pollard, K.S. & Kyrpides, N.C. New insights from uncultivated genomes of the global human gut microbiome. Nature 568, 505–510 (2019).

16. Pasolli, E. et al. Extensive Unexplored Human Microbiome Diversity Revealed by Over 150,000 Genomes from Metagenomes Spanning Age, Geography, and Lifestyle. Cell 176, 649–662 e620 (2019).

17. Friedman, J. & Alm, E.J. Inferring correlation networks from genomic survey data. PLoS Comput Biol 8, e1002687 (2012).

18. Sczyrba, A. et al. Critical Assessment of Metagenome Interpretation-a benchmark of metagenomics software. Nat Methods 14, 1063–1071 (2017).

19. Alneberg, J. et al. Binning metagenomic contigs by coverage and composition. Nat Methods 11, 1144–1146 (2014).

20. Backhed, F. et al. Dynamics and Stabilization of the Human Gut Microbiome during the First Year of Life. Cell Host Microbe 17, 852 (2015).

21. Kang, D.D., Froula, J., Egan, R. & Wang, Z. MetaBAT, an efficient tool for accurately reconstructing single genomes from complex microbial communities. PeerJ 3, e1165 (2015).

22. Jie, Z. et al. The vagino-cervical microbiome as a woman’s life history. *Under Revis*. Cell (2019).

23. Brito, I.L. et al. Transmission of human-associated microbiota along family and social networks. Nat Microbiol 4, 964–971 (2019).

24. Goltsman, D.S.A. et al. Metagenomic analysis with strain-level resolution reveals fine-scale variation in the human pregnancy microbiome. Genome Res 28, 1467–1480 (2018).

25. Heintz-Buschart, A. et al. Integrated multi-omics of the human gut microbiome in a case study of familial type 1 diabetes. Nat Microbiol 2, 16180 (2016).

26. Zeller, G. et al. Potential of fecal microbiota for early-stage detection of colorectal cancer. Mol Syst Biol 10, 766 (2014).

27. Nurk, S., Meleshko, D., Korobeynikov, A. & Pevzner, P.A. metaSPAdes: a new versatile metagenomic assembler. Genome Res 27, 824–834 (2017).

28. Bankevich, A. et al. SPAdes: a new genome assembly algorithm and its applications to single-cell sequencing. J Comput Biol 19, 455–477 (2012).

29. Bowers, R.M. et al. Minimum information about a single amplified genome (MISAG) and a metagenome-assembled genome (MIMAG) of bacteria and archaea. Nat Biotechnol 35, 725–731 (2017).

30. Parks, D.H., Imelfort, M., Skennerton, C.T., Hugenholtz, P. & Tyson, G.W. CheckM: assessing the quality of microbial genomes recovered from isolates, single cells, and metagenomes. Genome Res 25, 1043–1055 (2015).

31. Escapa, I.F. et al. New Insights into Human Nostril Microbiome from the Expanded Human Oral Microbiome Database (eHOMD): a Resource for the Microbiome of the Human Aerodigestive Tract. mSystems 3 (2018).

32. Forster, S.C. et al. A human gut bacterial genome and culture collection for improved metagenomic analyses. Nat Biotechnol 37, 186–192 (2019).

33. Zou, Y. et al. 1,520 reference genomes from cultivated human gut bacteria enable functional microbiome analyses. Nat Biotechnol 37, 179–185 (2019).

34. Liu, B. et al. Deep sequencing of the oral microbiome reveals signatures of periodontal disease. PLoS One 7, e37919 (2012).

35. Shi, B. et al. Dynamic changes in the subgingival microbiome and their potential for diagnosis and prognosis of periodontitis. MBio 6, e01926–01914 (2015).

36. Lassalle, F. et al. Oral microbiomes from hunter-gatherers and traditional farmers reveal shifts in commensal balance and pathogen load linked to diet. Mol Ecol 27, 182–195 (2018).

37. Schnorr, S.L. et al. Gut microbiome of the Hadza hunter-gatherers. Nat Commun 5, 3654 (2014).

38. Huerta-Cepas, J. et al. eggNOG 5.0: a hierarchical, functionally and phylogenetically annotated orthology resource based on 5090 organisms and 2502 viruses. Nucleic Acids Res 47, D309–D314 (2019).

39. Huerta-Cepas, J. et al. Fast Genome-Wide Functional Annotation through Orthology Assignment by eggNOG-Mapper. Mol Biol Evol 34, 2115–2122 (2017).

40. Zhang, C.C. et al. Proteomic analysis of Deinococcus radiodurans recovering from gamma-irradiation. Proteomics 5, 138–143 (2005).

41. Fredricks, D.N., Fiedler, T.L. & Marrazzo, J.M. Molecular identification of bacteria associated with bacterial vaginosis. N Engl J Med 353, 1899–1911 (2005).

42. Pan, H. et al. A gene catalogue of the Sprague-Dawley rat gut metagenome. Gigascience 7 (2018).

43. Pan, H. et al. A single bacterium restores the microbiome dysbiosis to protect bones from destruction in a rat model of rheumatoid arthritis. Microbiome 7, 107 (2019).

44. Nouioui, I. et al. Genome-Based Taxonomic Classification of the Phylum Actinobacteria. Front Microbiol 9, 2007 (2018).

45. Mark Welch, J.L., Rossetti, B.J., Rieken, C.W., Dewhirst, F.E. & Borisy, G.G. Biogeography of a human oral microbiome at the micron scale. Proc Natl Acad Sci U S A 113, E791–800 (2016).

46. Schmidt, T.S. et al. Extensive transmission of microbes along the gastrointestinal tract. Elife 8 (2019).

47. Feng, Q. et al. Gut microbiome development along the colorectal adenoma-carcinoma sequence. Nat Commun 6, 6528 (2015).

48. Xu, R., Wang, Q. & Li, L. A genome-wide systems analysis reveals strong link between colorectal cancer and trimethylamine N-oxide (TMAO), a gut microbial metabolite of dietary meat and fat. BMC Genomics 16 Suppl 7, S4 (2015).

49. Guertin, K.A., et al. Serum Trimethylamine N-oxide, Carnitine, Choline, and Betaine in Relation to Colorectal Cancer Risk in the Alpha Tocopherol, Beta Carotene Cancer Prevention Study. Cancer Epidemiol Biomarkers Prev 26, 945–952 (2017).

50. Wirbel, J. et al. Meta-analysis of fecal metagenomes reveals global microbial signatures that are specific for colorectal cancer. Nat Med 25, 679–689 (2019).

51. Cully, M. Microbiome therapeutics go small molecule. Nat Rev Drug Discov 18, 569–572 (2019).

52. Taylor, M.R. et al. Vancomycin relieves mycophenolate mofetil-induced gastrointestinal toxicity by eliminating gut bacterial beta-glucuronidase activity. Sci Adv 5, eaax2358 (2019).

53. Maier, L. et al. Extensive impact of non-antibiotic drugs on human gut bacteria. Nature 555, 623–628 (2018).

54. Blin, K. et al. antiSMASH 5.0: updates to the secondary metabolite genome mining pipeline. Nucleic Acids Research 47, W81–W87 (2019).

55. Shmakov, S. et al. Discovery and Functional Characterization of Diverse Class 2 CRISPR-Cas Systems. Mol Cell 60, 385–397 (2015).

56. Power, R.A., Parkhill, J. & de Oliveira, T. Microbial genome-wide association studies: lessons from human GWAS. Nat Rev Genet 18, 41–50 (2017).

57. Han, M. et al. A novel affordable reagent for room temperature storage and transport of fecal samples for metagenomic analyses. Microbiome 6, 43 (2018).

58. Fang, C. et al. Assessment of the cPAS-based BGISEQ-500 platform for metagenomic sequencing. Gigascience 7, 1–8 (2018).

59. Chen, S., Zhou, Y., Chen, Y. & Gu, J. fastp: an ultra-fast all-in-one FASTQ preprocessor. Bioinformatics 34, i884–i890 (2018).

60. Langmead, B. & Salzberg, S.L. Fast gapped-read alignment with Bowtie 2. Nat Methods 9, 357–359 (2012).

61. Shen, W., Le, S., Li, Y. & Hu, F. SeqKit: A Cross-Platform and Ultrafast Toolkit for FASTA/Q File Manipulation. PLoS One 11, e0163962 (2016).

62. Li, H. & Durbin, R. Fast and accurate short read alignment with Burrows-Wheeler transform. Bioinformatics 25, 1754–1760 (2009).

63. Torsten, S. barrnap 0.9: rapid ribosomal RNA prediction. https://github.com/tseemann/barrnap (2018).

64. Chan, P.P. & Lowe, T.M. tRNAscan-SE: Searching for tRNA Genes in Genomic Sequences. Methods Mol Biol 1962, 1–14 (2019).

65. Ondov, B.D. et al. Mash: fast genome and metagenome distance estimation using MinHash. Genome Biol 17, 132 (2016).

66. Mullner, D. fastcluster: Fast Hierarchical, Agglomerative Clustering Routines for R and Python. Journal of Statistical Software 53, 1–18 (2013).

67. Olm, M.R., Brown, C.T., Brooks, B. & Banfield, J.F. dRep: a tool for fast and accurate genomic comparisons that enables improved genome recovery from metagenomes through de-replication. Isme Journal 11, 2864–2868 (2017).

68. Pritchard, L., Glover, R.H., Humphris, S., Elphinstone, J.G. & Toth, I.K. Genomics and taxonomy in diagnostics for food security: soft-rotting enterobacterial plant pathogens. Analytical Methods 8, 12–24 (2016).

69. Segata, N., Bornigen, D., Morgan, X.C. & Huttenhower, C. PhyloPhlAn is a new method for improved phylogenetic and taxonomic placement of microbes. Nat Commun 4, 2304 (2013).

70. Hyatt, D., LoCascio, P.F., Hauser, L.J. & Uberbacher, E.C. Gene and translation initiation site prediction in metagenomic sequences. Bioinformatics 28, 2223–2230 (2012).

71. Buchfink, B., Xie, C. & Huson, D.H. Fast and sensitive protein alignment using DIAMOND. Nat Methods 12, 59–60 (2015).

72. Katoh, K. & Standley, D.M. MAFFT multiple sequence alignment software version 7: improvements in performance and usability. Mol Biol Evol 30, 772–780 (2013).

73. Capella-Gutierrez, S., Silla-Martinez, J.M. & Gabaldon, T. trimAl: a tool for automated alignment trimming in large-scale phylogenetic analyses. Bioinformatics 25, 1972–1973 (2009).

74. Nguyen, L.T., Schmidt, H.A., von Haeseler, A. & Minh, B.Q. IQ-TREE: a fast and effective stochastic algorithm for estimating maximum-likelihood phylogenies. Mol Biol Evol 32, 268–274 (2015).

75. Asnicar, F., Weingart, G., Tickle, T.L., Huttenhower, C. & Segata, N. Compact graphical representation of phylogenetic data and metadata with GraPhlAn. PeerJ 3, e1029 (2015).

76. Matsen, F.A., Kodner, R.B. & Armbrust, E.V. pplacer: linear time maximum-likelihood and Bayesian phylogenetic placement of sequences onto a fixed reference tree. Bmc Bioinformatics 11 (2010).

77. Price, M.N., Dehal, P.S. & Arkin, A.P. FastTree 2-Approximately Maximum-Likelihood Trees for Large Alignments. Plos One 5 (2010).

78. Eddy, S.R. Accelerated Profile HMM Searches. Plos Computational Biology 7 (2011).

79. Jain, C., Rodriguez-R, L.M., Phillippy, A.M., Konstantinidis, K.T. & Aluru, S. High throughput ANI analysis of 90K prokaryotic genomes reveals clear species boundaries. Nature Communications 9 (2018).

80. Ginestet, C. ggplot2: Elegant Graphics for Data Analysis. Journal of the Royal Statistical Society Series a-Statistics in Society 174, 245–245 (2011).

81. Bougeard, S. & Dray, S. Supervised Multiblock Analysis in R with the ade4 Package. Journal of Statistical Software 86, 1–17 (2018).

82. Oksanen, J. et al. vegan: Community Ecology Package. package version 2.5-2. https://CRAN.R-project.org/package=vegan (2018).

83. Kolde, R. pheatmap: Pretty Heatmaps. R package version 1.0.12. https://CRAN.R-project.org/package=pheatmap (2019).

84. Seemann, T. Prokka: rapid prokaryotic genome annotation. Bioinformatics 30, 2068–2069 (2014).

85. Scholz, M. et al. Strain-level microbial epidemiology and population genomics from shotgun metagenomics. Nature Methods 13, 435-+ (2016).

86. Morgan, X.C. et al. Dysfunction of the intestinal microbiome in inflammatory bowel disease and treatment. Genome Biology 13 (2012).

87. Kuhn, M. et al. caret: Classification and Regression Training. R package version 6.0-84. https://CRAN.R-project.org/package=caret (2019).

